# Neuraminidase activity mediates IL-6 production through TLR4 and p38/ERK MAPK signaling in MRL/lpr mesangial cells

**DOI:** 10.1101/2020.01.15.908160

**Authors:** Kamala P. Sundararaj, Jessalyn Rodgers, Peggi Angel, Bethany Wolf, Tamara K. Nowling

**Affiliations:** Department of Medicine, Division of Rheumatology; Medical University of South Carolina, Charleston, South Carolina; Department of Cell and Molecular Pharmacology and Experimental Therapeutics; Medical University of South Carolina, Charleston, South Carolina; Department of Public Health Sciences; Medical University of South Carolina, Charleston, South Carolina

**Keywords:** autoimmune disease, cytokine, interleukin 6 (IL-6), neuraminidase, toll-like receptor 4 (TLR4), mitogen-activated protein kinase (MAPK), cell signaling, mesangial cell

## Abstract

Mesangial cells (MCs), considered the immune cell of the kidney, secrete a number of cytokines including IL-6, which serves as an autocrine factor for MCs stimulating proliferation. IL-6 is associated with disease in patients and mouse strains with lupus nephritis, promoting tissue damage. Previously, we demonstrated the activity or levels of the enzyme neuraminidase (NEU) is increased in the kidneys of lupus mice and urine of human patients with nephritis and that NEU activity plays a role in mediating IL-6 secretion from lupus prone MRL/lpr primary mouse MCs. In this study, we further elucidate the mechanisms by which NEU activity mediates cytokine production by primary lupus prone MCs. MRL/lpr primary MCs were cultured with lupus serum to stimulate cytokine production in the absence or presence of NEU activity inhibitor. Our results show lupus serum increases NEU activity, and secretion of GM-CSF and MIP1α, in addition to IL-6, is significantly reduced when NEU activity is inhibited. mRNA expression of *Il-6* and *Gm-csf* was also increased in response to lupus serum, and reduced when NEU activity was inhibited. Using neutralizing antibodies to specific receptors, inhibitors of MAP kinase signaling pathways, and LPS stimulation we show TLR4 and p38/ERK MAPK play a role in NEU-mediated secretion of IL-6. Together, our results suggest NEU activity plays an important role in the response of lupus prone MCs to factor(s) in lupus serum that stimulates IL-6 expression and secretion through TLR4-p38/ERK MAPK signaling, likely through desialyation of one or more glycoproteins in this pathway.

## INTRODUCTION

Systemic lupus erythematosus (SLE) is an inflammatory, chronic autoimmune disorder involving multiple organs. Defects in the clearance of apoptotic cells lead to intracellular autoantigen release, autoantibody production, and immune complex deposition and inflammation in target organs. Glomerulonephritis is the leading cause of morbidity and mortality in lupus patients; however, the mediators of lupus nephritis (LN) remain incompletely known. Glycosphingolipid (GSL) metabolism is altered in lupus patients and lupus prone mice with nephritis compared to their non-nephritis counterparts and healthy controls (1), sharing similarities with other chronic kidney diseases shown to be mediated by GSL metabolism (2–5). However, the precise mechanisms by which GSL metabolism impacts chronic kidney diseases, including lupus nephritis, have not been elucidated. Proinflammatory cytokine IL-6 is upregulated in the kidney of mice and patients with lupus and was shown to be important for onset of nephritis in lupus mice (6–11). The altered GSL metabolism in MRL/lpr kidneys included increased activity of neuraminidase (1), which we demonstrated mediates IL-6 release from primary MRL/lpr mesangial cells (MCs) stimulated *in vitro* (12).

Neuraminidases (NEUs) (or sialidases) remove sialic acids from glycolipids and glycoproteins. There are four mammalian NEUs. NEU1 is typically located in the lysosome and was shown to be translocated to the plasma membrane of many cell types upon activation (13–16). NEU2 and NEU3 are located in the cytosol and plasma membrane, respectively (17,18), while NEU4 is found in the mitochondria and lysosome (19,20). NEUs also differ somewhat in substrate specificities and tissue distribution. NEU1 is ubiquitously expressed and is the major NEU expressed in the kidney (17,19,21–24). The only other NEU detectable in the kidney is NEU3 with message levels of Neu1 expressed approximately 40-fold higher than Neu3 (19) and NEU1 and NEU3 proteins are readily detected in the glomeruli of renal tissue sections of MRL/lpr lupus mice (12). While NEU1 was largely observed to desialylate glycoproteins, NEU3 showed a preference for gangliosides (25–29) and their activity at the plasma membrane impacts receptor activation and signaling in a variety of cell types (29–35).

To extend our previous studies and further elucidate the mechanisms by which NEU activity mediates MC function, we investigated the potential signaling pathways involved in NEU-mediated cytokine release in response to lupus serum. The activation of lupus prone MCs by soluble mediators present in lupus serum is not well established. Using lupus serum as a disease-relevant stimulator of MCs, we determined that NEU activity mediates the release of GM-CSF and MIP1α in addition to IL-6 by MRL/lpr lupus prone primary MCs. NEU activity likely mediates IL-6 release directly while effects on GM-CSF and MIP1α release are likely indirectly mediated by NEU activity. NEU mediated release of these cytokines occurs in part through TLR4-MAPK p38 or ERK signaling pathways in response to lupus serum and TLR4 ligand LPS that is likely due to removal of sialic acid residues from cell surface signaling proteins in this pathway by NEU.

## RESULTS

### Lupus serum stimulates cytokine secretion and mRNA expression in lupus prone MCs

To better define the effects of lupus serum in stimulating lupus prone MCs, Multiplex FlexMap3D bead array analysis was used to measure a panel of twenty cytokines. MRL/lpr MCs were cultured in the presence of serum taken from healthy non-autoimmune prone C57BL/6 (B6) mice or MRL/lpr nephritic mice. Secretion of seven cytokines (IL-6, IL-12, IL-17, MIP1α, GM-CSF, IFNγ, and IP-10) were increased by at least four-fold after addition of lupus serum and were decreased in the presence of the NEU inhibitor oseltamivir phosphate (OP). Individual ELISAs confirmed significant dose-dependent secretion of IL-6 (Fig. 1A, p=0.011) as we previously demonstrated (12), GM-CSF (Fig. 1B, p=0.013), and MIP1α (Fig. 1C, p<0.001) in response to lupus serum. We then analyzed the temporal release of these cytokines in response to 10% lupus serum stimulation. Media was collected from lupus MCs stimulated with lupus serum for 1, 3, and 6 hrs. Significantly increased secretion of all three cytokines was observed by 3 hrs following addition of lupus serum (Fig. 1D-F). Message levels of *Il-6* (Fig. 1G) and *Gm-csf* (Fig. 1H) were also significantly increased following lupus serum stimulation with significant increases in *Gm-csf* mRNA levels being observed within 1 hr of lupus serum stimulation (Fig. 1H). *Mip1*α mRNA levels were undetectable in unstimulated cells and remained at the threshold of minimal detection after stimulation making quantification unreliable (Fig. 1I). Thus, we were unable to conclude with any degree of certainty if *Mip1*α expression was increased following lupus serum stimulation.

**Fig. 1.**
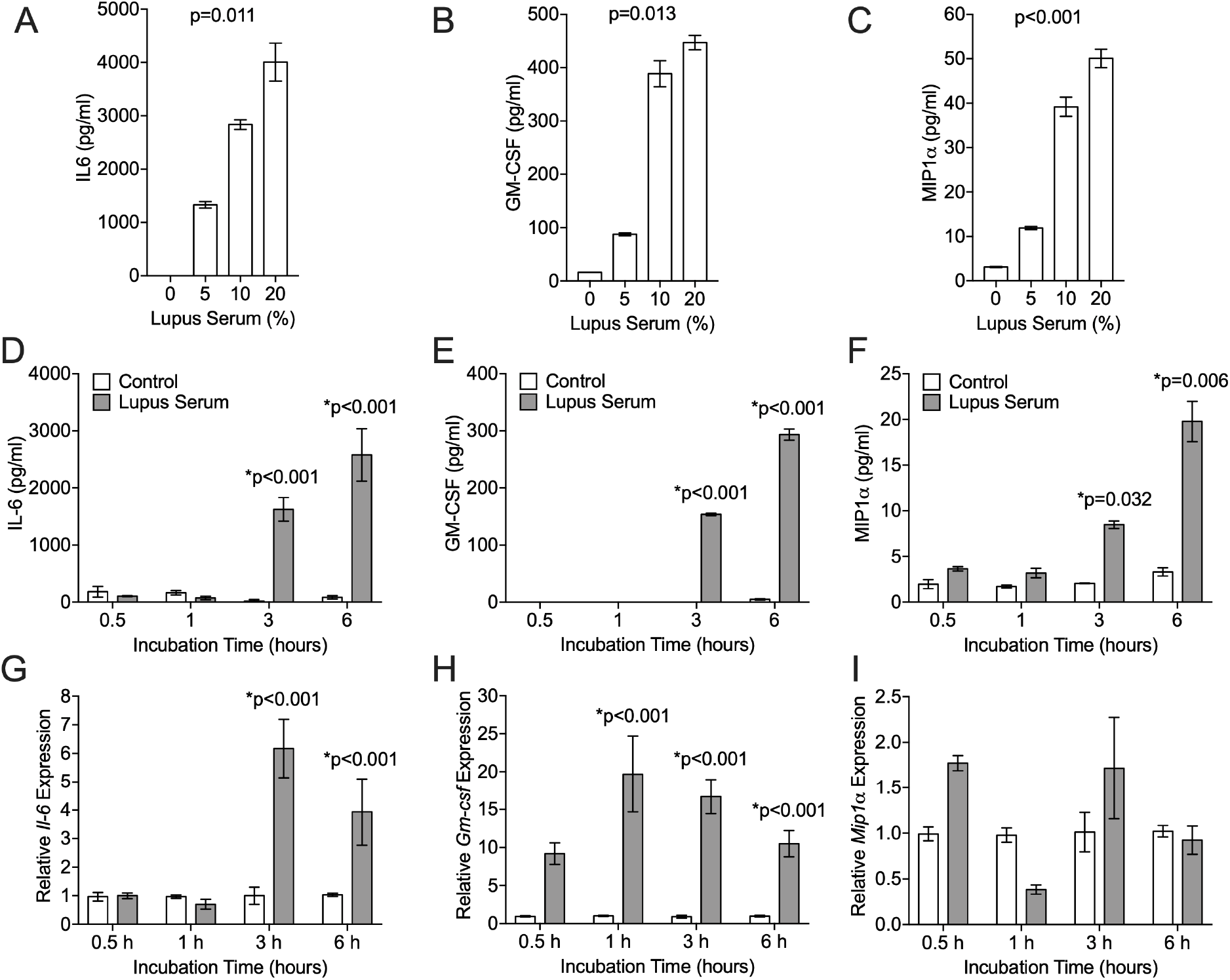
Lupus Serum stimulates expression and secretion of IL-6 and GM-CSF, and secretion of MIP1α from lupus prone primary MCs. IL-6 (A), GM-CSF (B), and MIP1α (C) were measured by individual ELISAs in the media of primary MRL/lpr MCs cultured for 6 hrs in the absence or presence of increasing concentrations of lupus serum. IL-6 (D), GM-CSF (E), and MIP1α (F) were measured by individual ELISAs in the media of primary MRL/lpr MCs cultured for the indicated time in the absence or presence of 10% lupus serum. *Il-6* (G), *Gm-csf* (H), and *Mip1α* (I) mRNA expression were measured by real-time RTPCR in primary MRL/lpr MCs cultured for 6 hrs in the absence or presence of 10% lupus serum. Experiments were performed at least three times with representative results presented.

### NEU activity mediates cytokine secretion from lupus serum stimulated primary lupus prone MCs

To confirm secretion of GM-CSF and MIP1α, in addition to IL-6 (12), in response to lupus serum are mediated by NEU activity, MCs were pretreated with OP or vehicle (water) and stimulated with lupus serum. This experiment confirmed all three cytokines were significantly decreased when MCs were pre-treated with OP prior to lupus serum stimulation (Fig. 2A-C). Interestingly, OP treatment blocked the increase in *Il-6* mRNA levels (Fig. 2D), but not the lupus serum-stimulated *Gm-csf* mRNA levels (Fig. 2E). In fact, pre-treating with OP further increased *Gm-csf* mRNA levels following lupus serum stimulation. A live-cell NEU activity assay demonstrated that lupus serum-stimulated NEU activity was returned to baseline activity levels by OP (Fig. 2F). These results suggest NEU activity mediates secretion of at least three cytokines by lupus prone MCs and that NEU activity may mediate IL-6 secretion at the message level in response to lupus serum.

**Fig. 2.**
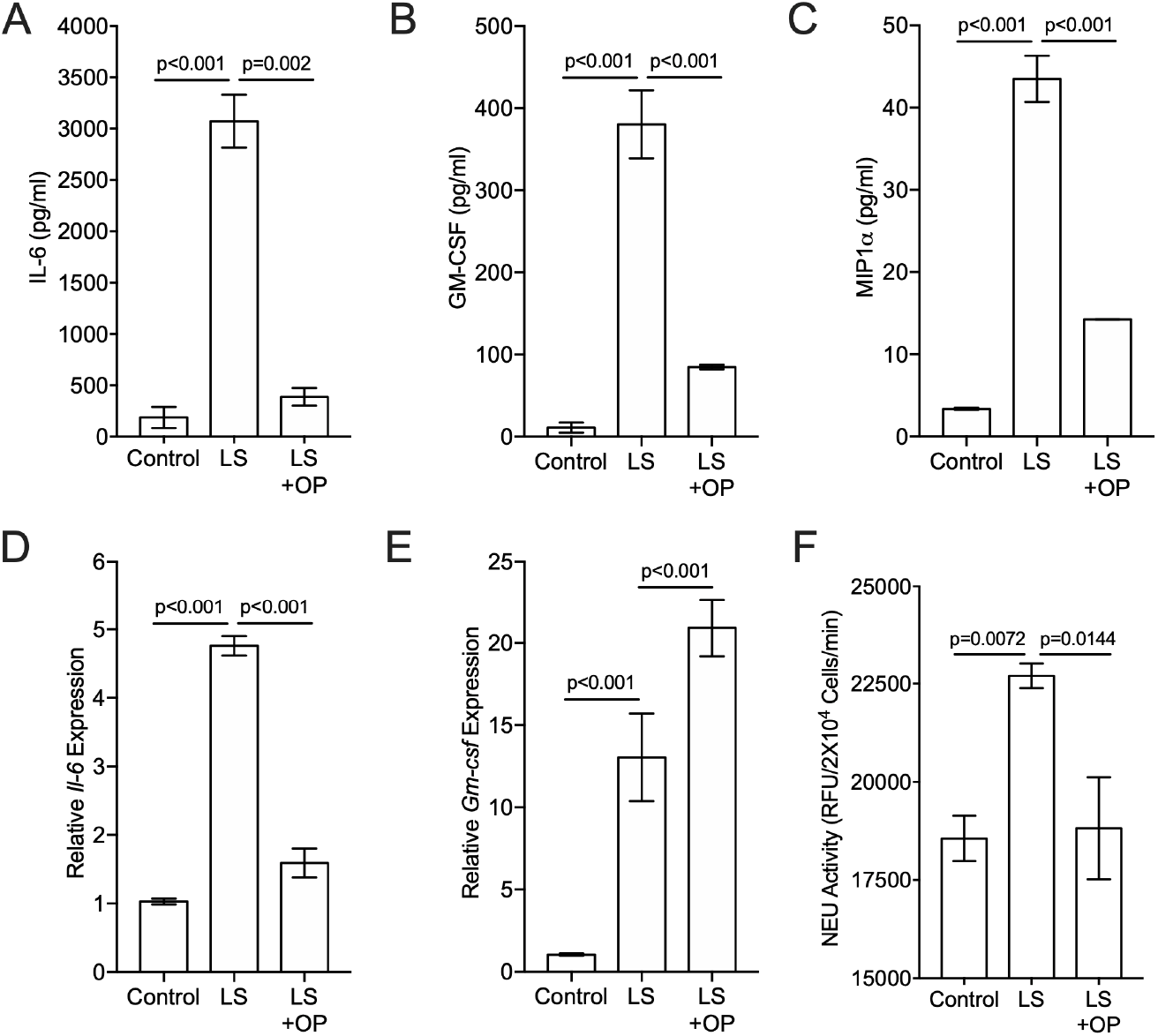
Inhibiting NEU activity significantly reduces lupus serum-stimulated secretion of IL-6, GM-CSF, and MIP1α, and expression of *Il-6*. Primary MRL/lpr MCs were cultured in the absence or presence of 10% lupus serum (LS) and NEU inhibitor oseltamivir phosphate (OP) as indicated in Materials and Methods. IL-6 (A), GM-CSF (B), and MIP1α (C) were measured by individual ELISAs in the media 6 hrs after addition of LS. *Il-6* (D) and *Gm-csf* (E) mRNA expression were measured by real-time RTPCR in the cells 6 hrs after addition of 10% LS. F) NEU activity was measured in cells after pretreatment with vehicle (water) or OP followed by stimulation with 10% LS for 3 hrs. NEU activity was measured after addition of NEU substrate to live cells. Experiments were performed at least three times with representative results presented.

### Lupus serum stimulates IL-6 secretion from MRL/lpr MCs through TLR4/MD2

The secretion of pro-inflammatory cytokines including IL-6, GM-CSF, and MIP1α can be directly stimulated through signaling pathways in MCs following ligand binding to cell surface proteins such as Fcγ receptors (FcγRs), toll like receptors (TLRs), annexin II (AnxA2), and TNF receptors (36–40). Therefore, the role of these proteins in lupus serum-stimulated primary MCs was investigated. MCs were incubated with neutralizing antibodies for cell-surface TLRs and FcγRs to block their interaction with putative ligands prior to addition of lupus serum. TLR4/MD2 neutralizing antibody significantly inhibited lupus serum-stimulated IL-6 secretion (Fig. 3A, p<0.001). Secretion of GM-CSF was slightly, but not significantly, reduced (Fig. 3B), and MIP1α was not significantly impacted (Fig. 3C) by anti-TLR4/MD2 antibodies. Antibodies to TLR4 only, AnxA2, TNFR, and isotype control IgGs also failed to reduce IL-6, GM-CSF, or MIP1α secretion (data not shown), indicating lupus serum stimulates IL-6 secretion from lupus prone MCs in part through TLR4 and MD2.

**Fig. 3.**
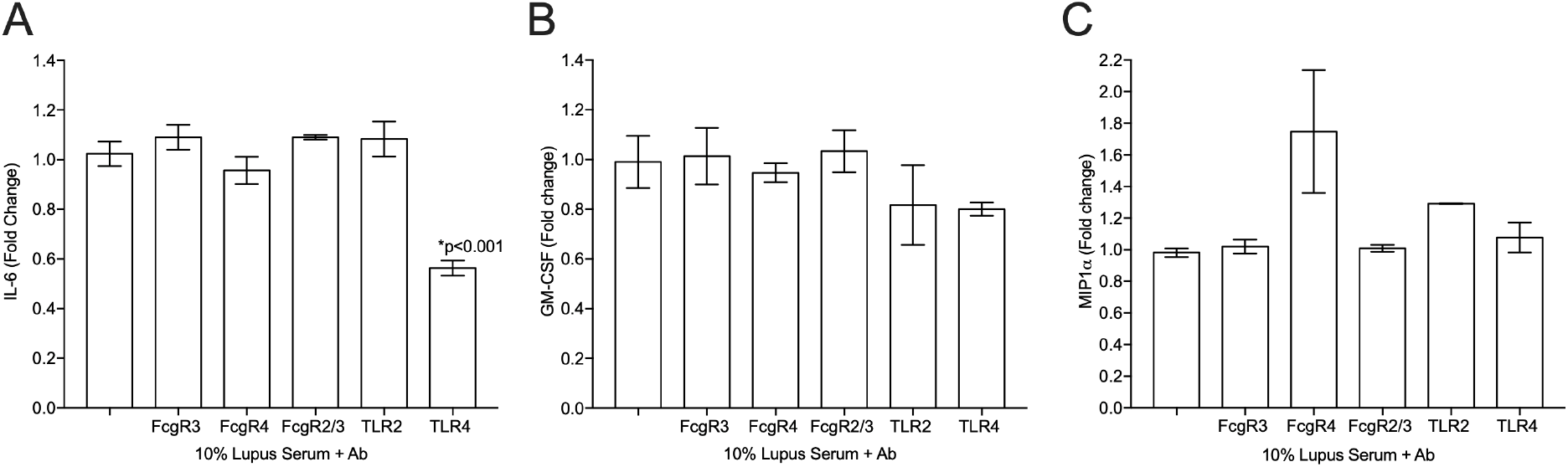
Blocking TLR4 significantly decreased lupus serum-stimulated IL-6 secretion. Primary MRL/lpr MCs were cultured with 10% lupus serum for 6 hrs following pretreatment without or with the indicated antibodies (Ab). IL-6 (A), GM-CSF (B), and MIP1α (C) were measured in the media by ELISA. Pretreatment with appropriate isotype controls failed to inhibit cytokine secretion (data not shown). Experiments were performed at least three times with representative results presented.

### NEU activity mediates IL-6 and GM-CSF secretion through TLR4 activation in MRL/lpr MCs

The TLR4 ligand LPS was then used to stimulate primary MRL/lpr MCs to examine if NEU activity mediates cytokine release specifically through a TLR4 pathway. LPS stimulated significant secretion of IL-6 (Fig. 4A, p=0.002), GM-CSF (Fig. 4B, p=0.013), and MIP1α (Fig. 4C, p=0.002). Increasing the concentration of LPS did not further increase the levels observed at 5 ng/ml indicating that release was maximal at this concentration. Thus, 5 ng/ml LPS was used to stimulate cells for all subsequent experiments. The secretion and mRNA expression of these cytokines by LPS stimulation followed a similar temporal trend as that observed following lupus serum stimulation. Significant increases in all three cytokines were observed in the media within 3 hrs of LPS addition (Fig. 4D-F). Message levels of *Il-6* (Fig. 4G) and *Gm-csf* (Fig. 4H) increased within 1 hr of stimulation, while *Mip1*α mRNA levels were essentially undetectable even after stimulation. Importantly, inhibiting NEU activity with the inhibitor OP, significantly reduced LPS-stimulated secretion of IL-6, GM-CSF, and MIP1α (Fig. 5A-C), and mRNA expression of *Il-6* and *Gm-csf* (Fig. 5D and E). Whereas *Gm-csf* mRNA levels were increased by OP treatment in lupus serum-stimulated MCs, they were decreased by OP in LPS-stimulated MCs.

**Fig. 4.**
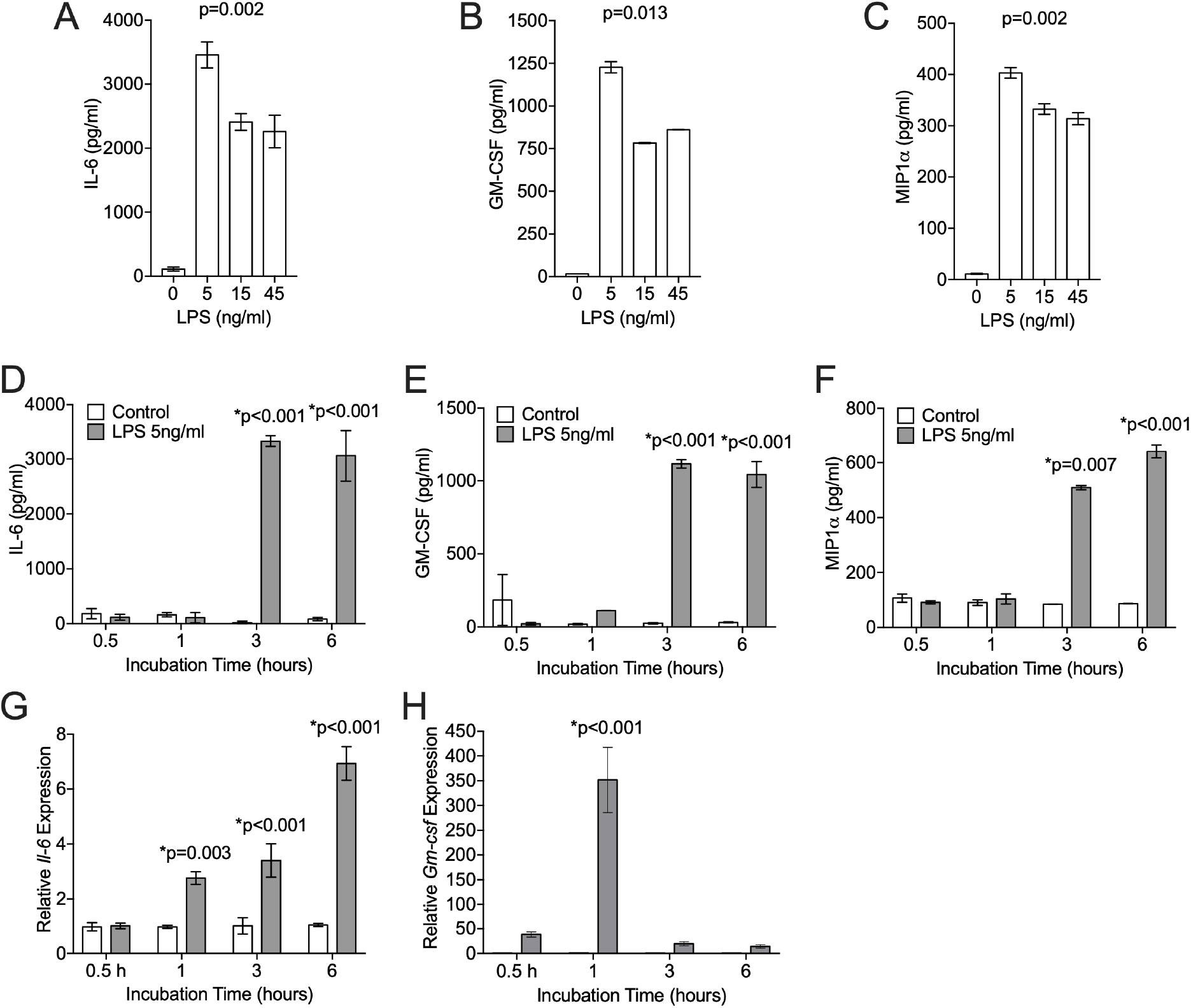
LPS stimulates expression and secretion of IL-6 and GM-CSF, and secretion of MIP1α from lupus prone primary MCs. IL-6 (A), GM-CSF (B), and MIP1α (C) were measured by individual ELISAs in the media of primary MRL/lpr MCs cultured for 6 hrs in the absence or presence of increasing amounts of LPS. IL-6 (D), GM-CSF (E), and MIP1α (F) were measured by individual ELISAs in the media of primary MRL/lpr MCs cultured for the indicated time in the absence or presence of 5 ng/ml LPS. *Il-6* (G) and *Gm-csf* (H) mRNA expression were measured by real-time RTPCR in primary MRL/lpr MCs cultured for 6 hrs in the absence or presence of 5 ng/ml LPS. Experiments were performed at least three times with representative results presented.

**Fig. 5.**
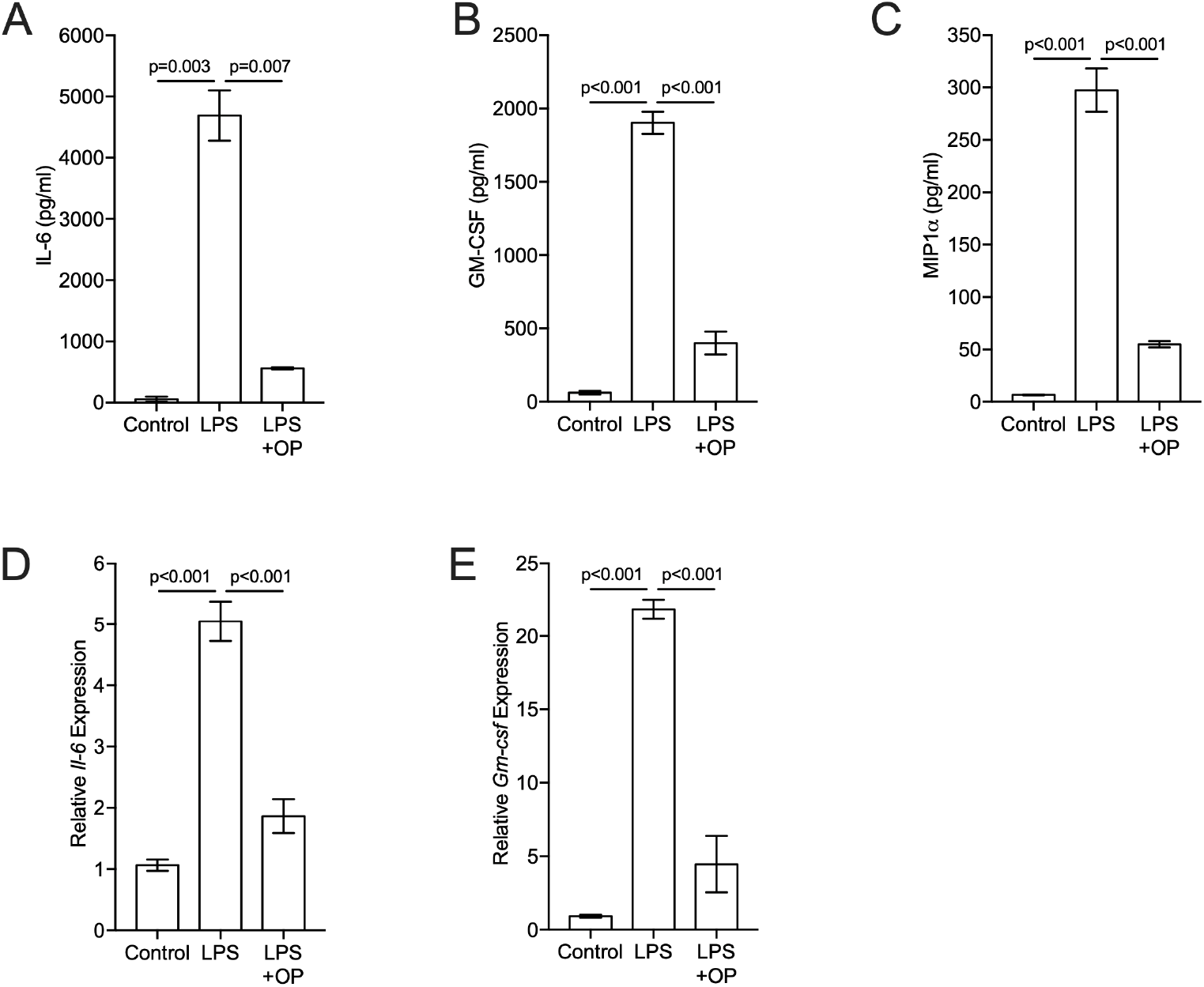
Inhibiting NEU activity significantly reduces LPS-stimulated secretion of IL-6, GM-CSF, and MIP1α, and expression of *Il-6* and *Gm-csf*. Primary MRL/lpr MCs were cultured in the absence or presence of 5 ng/ml LPS and NEU inhibitor oseltamivir phosphate (OP) as indicated in Materials and Methods. IL-6 (A), GM-CSF (B), and MIP1α (C) were measured by individual ELISAs in the media 6 hrs after addition of LPS. *Il-6* (D) and *Gm-csf* (E) mRNA expression were measured by real-time RTPCR in the cells 6 hrs after addition of LPS. Experiments were performed at least three times with representative results presented.

A live-cell NEU activity assay demonstrated significant NEU activity in response to LPS stimulation is significantly blocked by OP (Fig. 6A). These results suggest NEU activity mediates TLR4 signaling in upregulating expression and cellular release of IL-6 and GM-CSF in lupus prone MCs. To confirm that TLR4 is required for the LPS-stimulated release of IL-6, GM-CSF, and MIP1α, neutralizing antibodies were incubated with MCs prior to addition of LPS as performed with lupus serum. TLR4/MD2 neutralizing antibody significantly blocked LPS-stimulated IL-6 and GM-CSF (Fig. 6B) secretion. None of the neutralizing antibodies, including TLR4/MD2, had an effect on MIP1α secretion. Together, the lupus serum and LPS stimulation results suggest NEU-mediated secretion of IL-6 and GM-CSF from lupus prone MCs occurs in part through TLR4 activation, while NEU-mediated secretion of MIP1α occurs through a pathway that does not appear to require TLR2, TLR4, FcγRs, TNFRs, or AnxA2.

**Fig. 6.**
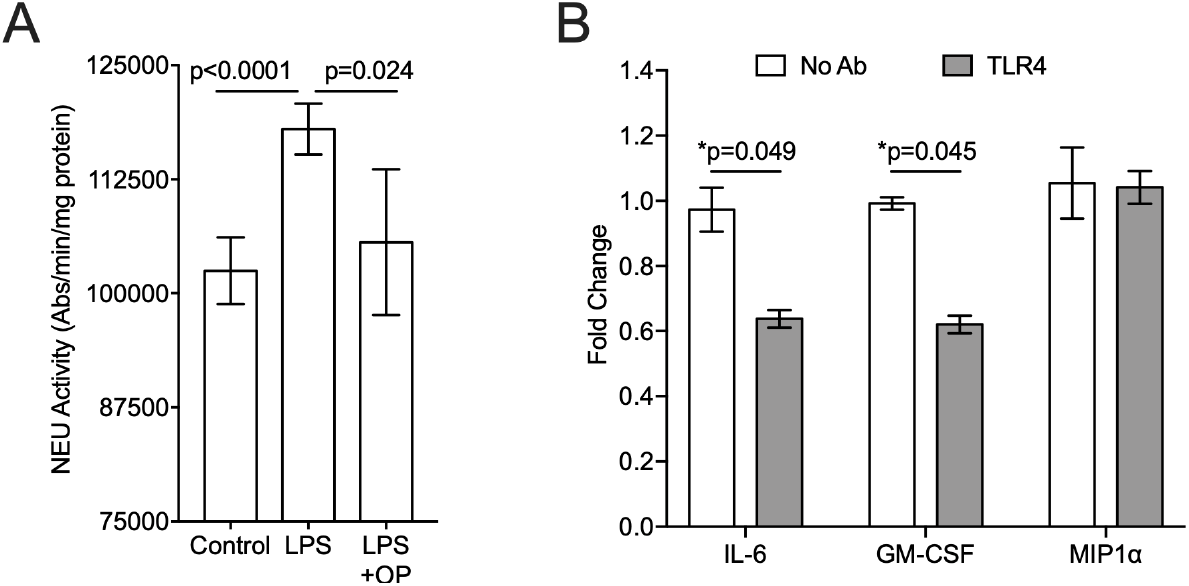
LPS stimulates NEU activity and LPS-stimulated cytokine release is decreased by anti-TLR4 antibodies. A) NEU activity was measured in primary MCs after pretreatment with vehicle (water) or OP followed by stimulation with 5 ng/ml LPS for 3 hrs. NEU activity was then measured after addition of NEU substrate to live cells. Experiments were performed at least three times with representative results presented. B) IL-6, GM-CSF, and MIP1α were measured in the media of primary MCs after pretreatment without (No Ab) or with TLR4 antibodies (Ab). Pretreatment with appropriate isotype control Abs failed to inhibit cytokine secretion or further increased cytokine secretion (data not shown). Experiments were performed three times and fold change relative to unstimulated cells averaged. Secretion in presence of LPS and No Ab was set to 1 for each cytokine and relative fold change in presence of TLR4 Ab presented.

### NEU activity mediates TLR4-MAPK signaling in secretion of IL-6 and GM-CSF from MRL/lpr MCs

Since the TLR4 receptor can signal through MAPK and PI3K pathways, we wanted to determine if MAPK (ERK, JNK, p38) or PI3K pathways play a role in NEU-mediated release of IL-6 in MCs. Inhibitors for each pathway were incubated with the cells prior to stimulation and included U0126, SB203580, and SP600125, inhibitors of ERK, p38, and JNK respectively, and wortmannin, an inhibitor of PI3K. Only ERK and p38 inhibitors significantly reduced IL-6 secretion from both lupus serum-stimulated (Fig. 7A, p<0.001 for both inhibitors) and LPS-stimulated (Fig. 7B, p<0.001 for both inhibitors) MCs. Inhibiting ERK also significantly reduced GM-CSF secretion from lupus serum-stimulated (Fig. 7A, p<0.001) and LPS-stimulated (Fig. 7B, p<0.001) MCs. None of the inhibitors reduced MIP1α secretion from MCs when stimulated with either lupus serum or LPS. Interestingly, JNK and PI3K inhibition significantly increased GM-CSF (p<0.001 for both) and PI3K inhibition significantly increased MIP1a secretion (p<0.001) in LPS-stimulated cells (Fig. 7B).

**Fig. 7.**
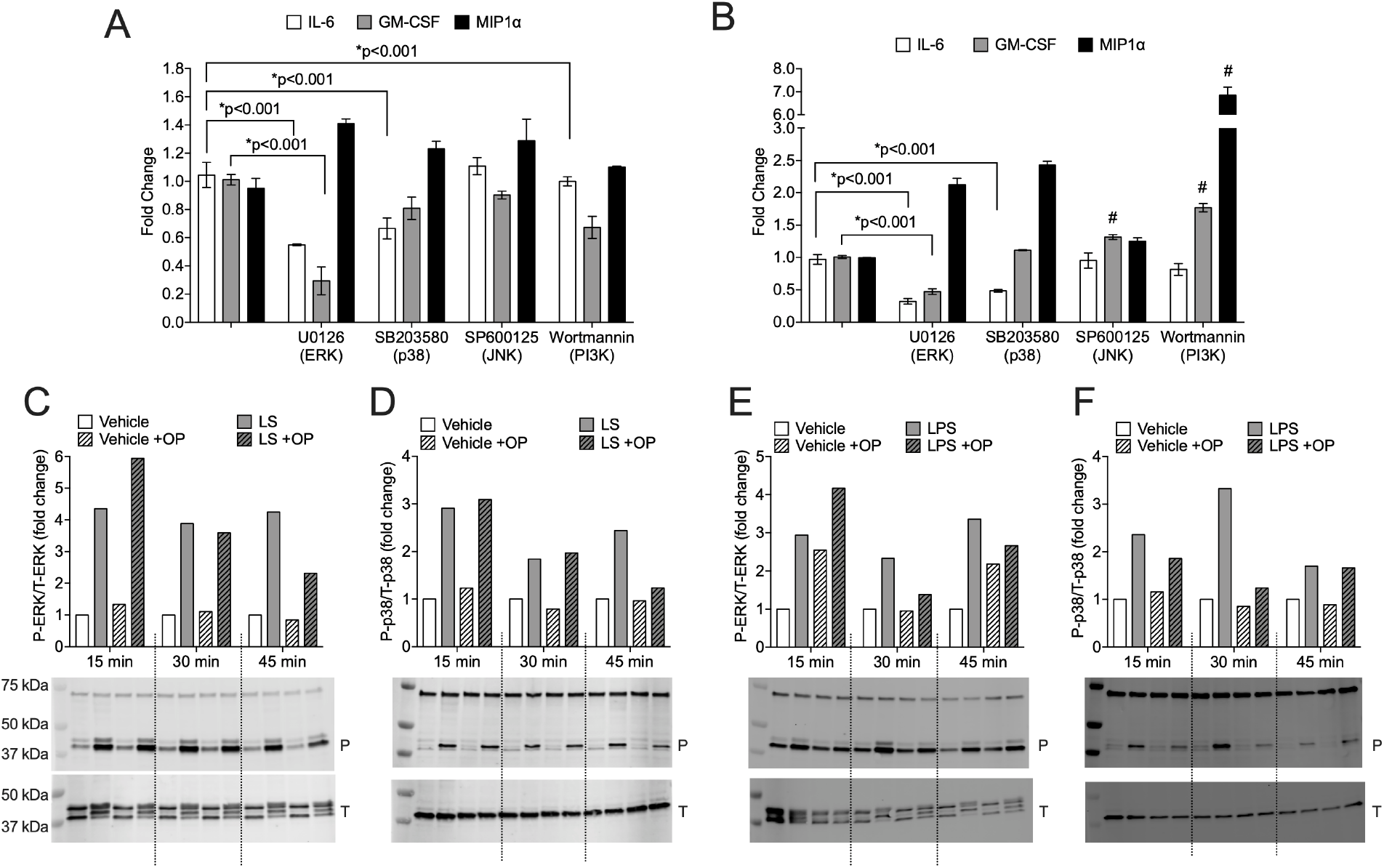
NEU activity mediates IL-6 and GM-CSF secretion through ERK or p38 MAPK signaling. MRL/lpr primary MCs were stimulated with 10% lupus serum (A, C, D) or 5 ng/ml LPS (B, E, F). Inhibitors for ERK (U0126), p38 (SB203580), JNK (SP600125), or PI3K (Wortmannin) were incubated with the cells prior to addition of 10% lupus serum (A) or 5 ng/ml LPS (B). Error bars represent standard deviation for the experiment presented. P-values represent significant changes of combined data from replicate experiments (see statistical analysis section). C-F) Cells were pretreated with vehicle (water) or NEU inhibitor oseltamivir phosphate (OP) and stimulated with 10% lupus serum (C and D) or 5 ng/ml LPS (E and F) for the indicated time. Whole cell extracts were prepared and subjected to western immunoblot using antibodies to phosphorylated ERK (P-ERK), total ERK (T-ERK), phosphorylated p38 (P-p38), and total p38 (T-p38). Ratio of P-ERK to T-ERK (C and E) and P-p38 to T-p38 (D and F) are graphed for the blots below each graph. P, phosphorylated p38 or ERK; and T, total p38 or ERK on the blots. Experiments were performed at least three times with representative results presented. #, p<0.001

To demonstrate NEU activity plays a role in ERK and p38 MAPK signaling in stimulated MCs, MRL/lpr primary MCs were incubated with OP followed by stimulation with lupus serum for 15, 30, and 45 min. Within 15 min of lupus serum stimulation, activation of both ERK (Fig. 7C) and p38 (Fig. 7D) was increased as indicated by the increase of their phosphorylated forms. The increases in phosphorylated ERK and p38 were blocked by the NEU inhibitor OP at 45 min, indicating NEU activity mediates ERK and p38 MAPK signaling in response to lupus serum. This experiment was repeated in LPS-stimulated MCs to examine the role of NEU activity in ERK and p38 MAPK signaling specifically through TLR4 activation. Similar to the lupus serum-stimulated MCs, phosphorylation of ERK (Fig. 7E) and p38 (Fig. 7F) was observed within 15 min of LPS stimulation, and inhibiting NEU activity with OP reduced the levels of phosphorylated forms of ERK and p38 at 30 min. Depending on the lupus serum lot used for stimulation, the observed reduction in levels in the presence of OP was consistent at 45 min for lupus serum-stimulated and at 30 min for LPS-stimulated MCs. Together, these results suggest NEU mediates TLR4-ERK/p38 MAPK IL-6 and GM-CSF cytokine release in lupus serum-stimulated lupus MCs.

### Inhibiting NEU activity significantly increases sialic acid-containing glycoproteins in lupus serum-stimulated MCs

Sialic acid-containing glycolipids and glycoproteins serve as substrates of NEUs and sialylation plays an important role in TLR4 activation and signaling in other cell types (41–43). We recently showed that sialic acid-containing glycosphingolipid GM3 is significantly increased in lupus serum-stimulated MCs when NEU activity is blocked (manuscript under review). Here, we profiled N-glycans to determine if blocking NEU activity also increases sialic acid-containing N-linked glycoproteins (N-glycans) in lupus-serum stimulated MCs in the absence or presence of OP. Fig. 8 presents two N-glycans that exhibited significantly increased sialylation in response to OP. Sialylation of N-glycan m/z 2122 (Fig. 8A) and m/z 2853 (Fig. 8B) were significantly increased in response to lupus serum, which were further significantly increased when NEU activity was inhibited. These results suggest NEU may mediate cytokine release by desialylating glycoproteins or glycolipids in signaling pathways leading to cytokine secretion in lupus serum-stimulated MRL/lpr MCs.

**Fig. 8.**
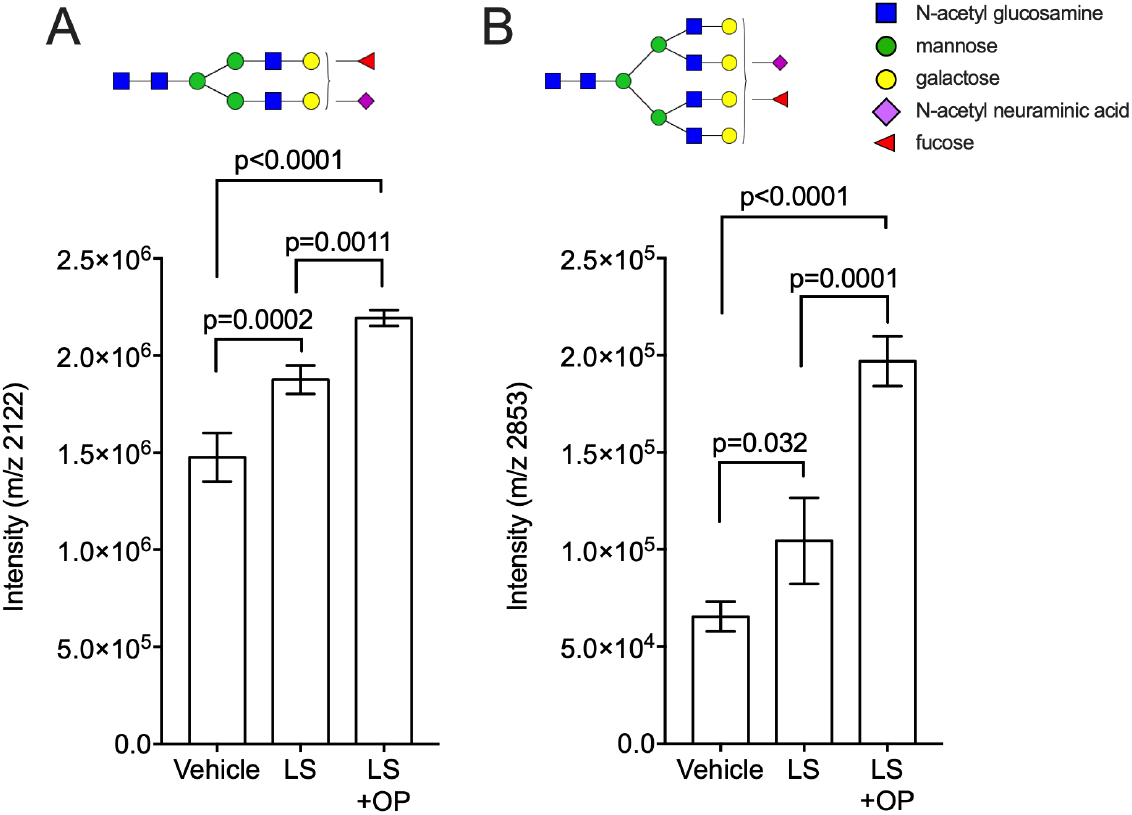
Blocking NEU activity significantly increased sialic acid-containing glycoproteins in lupus serum-stimulated MCs. MRL/lpr Primary MCs were pretreated with vehicle (water) or the NEU activity inhibitor oseltamivir phosphate (OP) and stimulated with 10% lupus serum for 6 hrs. N-glycans were analyzed by MALDI-FTICR. Two sialic acid-containing N-glycans with significant increases in sialylation are presented: m/z 2122 (A) and m/z 2853 (B). Vehicle, MCs received water only. Lupus serum, MCs received vehicle and 10% lupus serum. Lupus serum +OP, MCs received OP and 10% lupus serum.

## DISCUSSION

One of the earliest events in the pathogenesis of lupus nephritis is the deposition of immune complexes to promote cytokine production by mesangial cells (MCs), as well as podocytes. Cytokines, including IL-6, released by these resident renal cells promote the infiltration of immune cells, further stimulation of cytokine production by and proliferation of MCs, and subsequent tissue damage. Blocking or reducing IL-6 in lupus prone mice improved disease (7,10,11) and highlights the important role IL-6 plays in the progression of lupus. However, anti-IL-6 treatment of lupus patients with class III or IV nephritis in a small clinical trial was not efficacious (44). A recent study demonstrated IL-6 is specifically produced by glomerular MCs in NZM2328 lupus prone mice with glomerulonephritis (GN) (45). Delineating the precise mechanisms involved in IL-6 signaling in MCs will identify additional targets for reducing IL-6 levels as potential therapeutic interventions in LN. Our previously published results (12) and the data presented here indicate neuraminidase (NEU) activity plays an important role in mediating IL-6 production in MRL/lpr lupus prone MCs *in vitro*.

We previously demonstrated NEU activity mediates the release of IL-6, but not MCP-1, from lupus prone MRL/lpr MCs following stimulation with an immune complex mimic (heat-aggregated IgG) and lupus serum (12). Here we extended those studies and demonstrated IL-6 secretion by lupus serum-stimulated MCs is likely directly mediated by NEU activity through TLR4 and MAPK p38 or ERK signaling since both IL-6 secretion and message levels were significantly decreased when NEU activity was inhibited. The pro-inflammatory activity of lupus serum is largely thought to require the presence of circulating immune complexes. The presence of autoantibodies and immune complexes, with the ability to activate FcγRs and TLRs, leads to increased production of inflammatory mediators, such as cytokines, chemokines, and reactive oxygen species. In our studies, FcγRs do not appear to be required in lupus serum-stimulated secretion of IL-6, GM-CSF or MIP1α in MRL/lpr MCs. These results are in agreement with previous studies demonstrating that MRL/lpr mice lacking the Fcγ chain still develop renal disease (46), and FcγRs are dispensable for activation of MRL/lpr MCs by pathogenic anti-DNA antibodies (36). Removing IgG complexes from lupus serum using anti-IgG beads did not reduce IL-6 secretion (unpublished data) suggesting lupus serum-stimulated IL-6 secretion by MCs does not require IgG immune complexes either. Interestingly, lupus serum heated to 56°C for 30 min failed to stimulate secretion of IL-6 in MRL/lpr MCs (unpublished data), suggesting complement may play a role. Future studies are aimed at determining if complement in lupus serum plays a role in NEU-mediated secretion of IL-6 by MCs.

NEU activity can mediate cell signaling by removing sialic acids (desialylating) from lipids or proteins. Gangliosides, sialic acid-containing GSLs, present in cell membranes can modulate cell functions by organizing lipid rafts to promote receptor-mediated signal transduction (47). Increasing GM3, the simplest sialic acid-containing ganglioside from which all other gangliosides are generated, was shown to prevent MC activation and proliferation (48). Treating lupus serum-stimulated MCs with OP to block NEU activity resulted in a significant increase of GM3 levels (manuscript under review) and we show here that this treatment also results in a significant decrease in IL-6 production. Thus, GM3 may play a role in the signaling pathway for IL-6 production in lupus-serum stimulated MCs.

Alternatively or additionally, the function of NEU in mediating IL-6 production may be through the desialylation of glycoproteins involved in the IL-6 signaling pathway. We observed small, yet significant increases in sialylated N-glycans when the MRL/lpr MCs were stimulated with lupus serum, which were further highly increased in the presence of OP. Sialic acids (SAs) can mask receptors from ligands to prevent signaling (49). In particular, TLR4, which we demonstrated is necessary in both lupus serum- and LPS-stimulated IL-6 production in MCs, can be activated by NEU. TLR4 diffuses into lipid rafts upon LPS stimulation to propagate signaling of gene expression (50) and removal of SAs by NEU enhanced binding and subsequent signaling through TLR4 (31,43,51,52). Increasing sialylation *in vivo* by treating mice with exogenous SAs reduced TLR4 activation and neutralized LPS toxicity-induced renal injury (53). Optimal TLR4 signaling requires MD2, both of which contain multiple N-glycans required for function that can be sialylated. TLR4 activation in other cell types was shown to trigger NEU1 translocation to the cell surface to desialylate TLR4 and MD2 to enhance TLR4 dimerization and allow signaling (42,43,54). Although TLR4 was not readily detected on MCs from healthy non-autoimmune mice, it was observed on MCs from MRL/lpr lupus mice (37) and TLR4 deficiency in another lupus prone mouse strain decreased renal disease (55). Our N-glycan results demonstrate OP treatment increases sialylation of N-glycans, paralleling significant reduction in IL-6 secretion in lupus serum-stimulated MCs. The N-glycan structures of TLR4 are unknown so we were unable to determine if the N-glycans identified in our analysis belong to TLR4. Future studies are focused on determining if TLR4 is a target of NEU and is desialylated in lupus MCs.

NEU activity also mediated GM-CSF and MIP1α secretion from lupus prone MCs. Blocking TLR4 in lupus serum-stimulated cells resulted in a slight reduction in GM-CSF secretion and blocking TLR4 in LPS-stimulated cells to isolate the TLR4 pathway resulted in a significant reduction of GM-CSF. Furthermore, blocking NEU activity significantly reduced *Gm-csf* message levels in LPS- and not lupus serum-stimulated cells. Together, these data suggest TLR4 may play a role, but other receptors are also likely involved in NEU-mediated GM-CSF secretion in response to lupus serum. MIP1α was not decreased in OP-treated cells when TLR4 or MAPK signaling was blocked, indicating NEU activity may mediate MIP1α secretion indirectly or through another mechanism.

In summary, we demonstrated stimulation of lupus prone primary MCs with lupus serum or LPS results in the secretion of IL-6, GM-CSF, and MIP1α and is mediated in part by NEU activity. We further showed NEU-mediated IL-6 production likely occurs through p38/ERK MAPK signaling following TLR4 activation. Glycosphingolipid and N-glycan analyses suggest NEU activity may mediate IL-6 production by directly desialylating glycolipids and/or glycoproteins to enhance TLR4-p38/ERK signaling in response to lupus serum. Together, our results suggest removal of SAs by NEU may regulate lupus serum-stimulated cytokine release by mediating TLR4-MAPK signaling in lupus prone MCs.

## EXPERIMENTAL PROCEDURES

### Reagents

Antibodies used include anti-FcγR3 and anti-TNFR1/TNFRSF1A (R&D Systems, Minneapolis, MN); anti-FcγR2/3, anti-FcγR4, anti-TLR2, and anti-TLR4/MD2 (BioLegend, San Diego, CA); anti-TNFR2 (BioXCell, West Lebanon, NH); anti-Annexin A2 (Abcam, Cambridge, MA); and isotype controls hamster IgG and mouse IgG (BioLegend, San Diego, CA), and rat IgG (Rockland Immunochemicals, Limerick, PA). Inhibitors used include: oseltamivir phosphate (Santa Cruz Biotechnology, Dallas, TX); ERK MAPK inhibitor U0126, JNK MAPK inhibitor SP600125, and PI3K inhibitor wortmannin (Calbiochem, San Diego, CA); p38 MAPK inhibitor SB203580 (AdipoGen, San Diego, CA). Cell stimulants include: Ultra-pure Lipopolysaccaride (LPS) (Invivogen, San Diego, CA); Lupus serum was obtained from 16-18 week-old MRL/lpr mice and pooled. Each pooled serum collection was tested for endotoxin using Pierce Chromogenic Endotoxin Quantification Kit (Thermo Scientific, Rockford, IL) and each “batch” used for experiments had undetectable levels of endotoxin.

### Generation and culturing of primary mesangial cell lines

Kidney glomeruli were isolated from 6-week-old pre-nephritic female MRL/lpr mice for generation of two independent primary mesangial cell lines (MCs) as described previously (12). Each primary MC line was characterized by flow cytometry at passage 5, by IHC at passage 6, and by RT-PCR at passages 5–9 as performed previously (56) and were 93% positive for αSMA by flow cytometry. All cell lines tested negative for mycoplasma (Applied StemCell, Inc, Milpitas, CA). Cells were grown in a 5% CO_2_ atmosphere at 37°C in MEM/D-valine medium with 20% fetal bovine serum, 1% penicillin/streptomycin and ITS supplement (5 μg/ml insulin, 5 μg/ml transferrin, 5 ng/ml sodium selenite) (GenDEPOT, Houston, TX; Sigma-Aldrich, St. Louis, MO) and cell passages 6-9 were utilized for all experiments. Cells were pretreated with 500 μM OP for 16 hrs, 40 μg/ml antibodies for 2 hrs, and 10 mM MAPK/PI3K inhibitors for 2 hrs prior to stimulation with 5-20% lupus serum or 5-45 ng/ml LPS for 0.5-24 hrs as indicated in the figures or figure legends. Results from each line showed similar trends with data presented from one line.

### Neuraminidase activity assay

Primary MCs (20,000 cells per well) were grown in a CoStar 3603 black 96-well clear-bottom plate, pre-treated with 500 μM OP and stimulated with 10% lupus serum or 5ng/ml LPS for 3 hrs. After incubation, media was collected, and cells were washed with 37°C warmed sterile filtered Tris-buffered saline (TBS) pH 7.4. Neuraminidase activity was measured using a SpectraMax i3c fluorometer with SoftMax Pro 6 software (Molecular Devices, San Hose, CA). Cells were incubated with 15 μM substrate 2′ (4-methylumbelliferyl)-α-D-*N*-acetylneuraminic acid (4MU-NANA) in 37°C warmed sterile filtered TBS (Sigma Aldrich; St. Louis, MO). Fluorescence intensity (excitation: 365 nm, emission: 460 nm) was measured at baseline prior to substrate addition and in the presence of substrate. Relative fluorescence units (RFU) per 2×10^4^ cells per min incubation with substrate is presented on the graph. Total protein content was consistent between untreated and treated cells. Significant differences were determined by one-way ANOVA with Tukey’s multiple comparisons test. Adjusted p-values are presented on the graphs.

### FlexMap 3D and ELISA analyses

Cytokines and chemokines analyzed in media of primary MC cultures by multiplex bead array analysis used a FlexMap3D platform (Luminex, Austin, TX) and Cytokine Mouse Magnetic 20-Plex Panel kit (Invitrogen/ThermoFisher Scientific, Waltham, MA). The FlexMap3D results were confirmed by individual commercially available enzyme-linked immunoassay (ELISA) detection kits. ELISAs were used to measure IL-6 (Biolegend, San Diego, CA), GM-CSF (R&D systems, Minneapolis, MN), and MIP1α (R&D systems, Minneapolis, MN) in primary MC supernatants according to the manufacturer’s instructions. Protein concentration of cells was measured using a Micro BCA Protein Assay Kit (Thermo Scientific, Rockford, IL) and used to normalize cytokine levels with final concentrations adjusted to an equivalent of 25 μg of protein for comparison across experiments.

### Real-time PCR

Total RNA was extracted from primary MCs using the RNeasy kit (Qiagen, Hilden, Germany) according to the manufacturer’s instructions, and cDNA was reversely transcribed with 0.5–1 μg RNA using the iScript cDNA synthesis kit as described in the kit (Biorad, Hercules, CA). The gene expression of *Il-6*, *Gm-csf*, *Mip1*α and housekeeping gene β*-actin* was evaluated by quantitative RT-PCR using the LightCycler 480 SYBR Green 1 Master kit and LightCycler 480 II (Roche, Indianapolis, IN) and oligos: *mIl-6* Forward 5’-TAGTCCTTCCTACCCCAATTTCC-3’ and Reverse 5’-TTGGTCCTTAGCCACTCCTTC-3’; *mGm-csf* Forward 5’-GGCCTTGGAAGCATGTAGAGG-3’ and Reverse 5’-GGAGAACTCGTTAGAGACGACTT-3’; *mMip1*α Forward 5’-ACTGCCTGCTGCTTCTCCTACA-3’ and Reverse 5’-AGGAAAATGACACCTGGCTGG-3’; *m*β-actin 5’-AGATTACTGCTCTGGCTCCTAG-3’ and Reverse 5’-CCTGCTTGCTGATCCACATC-3’. The PCR mixture was heated initially at 95 °C for 5 min, followed by 40 cycles of denaturation at 95 °C for 10 s and combined annealing/extension at 60 °C for 30 s. All reactions were performed in triplicate and normalized to β*-actin*. Gene expression data analysis was performed by using the comparative threshold cycle (CT) method.

### Immunoblotting

Mesangial cell extracts were prepared by incubating cells with RIPA buffer (50 mM Tris-HCl pH 7.5, 150 mM NaCl, 1% Triton X-100, 0.5% NaDeoxycholate, 0.1% SDS, 25 mM DTT, 10 mM EDTA, 1 mM PMSF). After centrifugation at 4°C for 10 minutes at 14,000 rpm, the supernatant was collected and protein concentration was determined using a micro BCA assay (Pierce/ThermoScientific, Rockford, IL). Samples (20–40 μg per well) were electrophoresed in a 4-20% Criterion TGX gel (Biorad, Hercules, CA), and transferred to a polyvinylidene difluoride membrane. Total and phosphorylated MAPK were immunoblotted with anti-mouse total and anti-mouse phosphorylated MAPK primary antibodies (Cell Signaling Technology, Danvers, MA), respectively. Detection was further enhanced with an anti-rabbit biotin antibody (Thermo Scientific, Rockford, IL) and AlexaFluor-647 streptavidin-conjugated antibody (Life Technologies, Grand Island, NY). Blots were scanned using Odyssey Infrared Imaging system and software (LI-COR, Lincoln, NE) and band density was quantified using NIH ImageJ. The results are presented as the ratios of phosphorylated MAPK to total MAPK.

### N-glycan analysis

Glycoprotein N-glycans were measured in intact cells grown on glass slides and cultured with lupus serum in the absence or presence of OP as described above. Cells were incubated with OP for 16 hrs and media replaced containing no lupus serum, 10% lupus serum, or 10% heat-treated lupus serum for 3 hrs. Heat-treated lupus serum was incubated at 56 °C for 30 min. Heat-treated lupus serum did not stimulate IL-6 secretion (data not shown). Following stimulation, cells were prepared for N-glycan profiling as described previously (57). Briefly, the array was washed in cold PBS and fixed in neutral buffered formalin followed by delipidation using Carnoy’s Solution (10% glacial acetic acid, 30% chloroform, 60% 200 proof ethanol) for three minutes. Deglycosylation was performed by spraying each array with PNGase F Prime (N-zyme Scientifics) using an M3 TM-Sprayer (HTX Technologies) with 10 passes at 25 μL/min, 1200 mm/s, 45°C, 3 mm spacing between passes with 10 psi nitrogen gas and incubated for 2 hours at 37°C and ≥80% relative humidity. MALDI matrix α-cyano-4-hydroxycinnamic acid (CHCA, Sigma; 50% acetonitrile/0.1% trifluoroacetic acid) was sprayed onto the cells with an M3 TM-Sprayer (HTX Technologies) using 10 passes at 70 μL/min, 1300 mm/s, 79°C, 2.5 mm spacing between passes and 10 psi nitrogen gas. Two passes of ammonium phosphate monobasic (5 mM) were applied using same parameters as for matrix. N-glycans were profiled using a Fourier Transform Ion Cyclotron Resonance Mass Spectrometer (FT-ICR; 7 Tesla solariX^™^, Bruker Scientific, LLC) equipped with a MALDI source. Transients of 512 kiloword were acquired in broadband positive ion mode over m/z 500-5,000 with a calculated on-tissue mass resolution at full width half maximum of 81,000 at m/z 1400. Each pixel consisted of 1200 laser shots rastered over a 500 μm diameter area. A lockmass on primary N-glycan peak m/z 1663.5814 (composition Hex5HexNAc4 + 1Na) was used to maintain mass accuracy during acquisition.

### Statistical analyses

Association between treatment or duration of exposure to lupus serum or LPS in the absence or presence of OP with outcomes (e.g. protein, mRNA levels, GSLs) were evaluated using a linear mixed model approach. Models with one treatment variable included a fixed effect for treatment and models that included treatment and time included fixed effects for treatment, time, and the treatment X time interactions. All experiments were performed at least two times with similar results, data were combined for the statistical analyses, and representative graphs are presented. Variability in extent of cytokine production was observed from experiment to experiment due to differences in passage number of the primary MCs used or batches of pooled serum in the lupus serum-stimulated cells. Therefore, all models included a random batch effect to account for correlation between experimental results from the same batch. Model assumptions were checked graphically and transformations were considered as needed. Pairwise differences between groups were evaluated using linear contrasts and a Bonferroni correction was applied to adjust for multiple comparisons. All analyses were conducted in SAS v. 9.4 (SAS Institute, Cary, NC). Global p-values are provided for analyzing lupus serum or LPS dose response of cytokine production. Significant differences in the glycan analyses were determined by one-way ANOVA with Tukey’s multiple comparisons test. Adjusted p-values are presented on all graphs.

## ACKNOWLEDGEMENTS

The authors thank Ivan Molano for assistance in performing the FlexMap 3D assay.^1^

## Footnotes

1 This work was supported by: 1) Office of the Assistant Secretary of Defense for Health Affairs through the Peer-Reviewed Medical Research Program Lupus Topic Area Award W81XWH-16-1-0640 (funding awarded to T. K. Nowling); 2) NIH Clinical Center grant U01 CA242096 and NIH Exploratory Center grant P20 GM103542 awarded to the South Carolina Center of Biomedical Research Excellence (COBRE) in Oxidants, Redox Balance, and Stress Signaling (supporting P.M. Angel). Opinions, interpretations, conclusions, and recommendations are those of the authors and are not necessarily endorsed by the Department of Defense. The content is solely the responsibility of the authors and does not necessarily represent the official views of the National Institutes of Health.

